# Integrated single cell functional-proteomic profiling of human skeletal muscle reveals a shift in cellular specificity in nemaline myopathy

**DOI:** 10.1101/2024.10.17.618209

**Authors:** Robert A.E. Seaborne, Roger Moreno-Justicia, Jenni Laitila, Chris T. A. Lewis, Lola Savoure, Edmar Zanoteli, Michael W Lawlor, Heinz Jungbluth, Atul S. Deshmukh, Julien Ochala

**Affiliations:** Centre of Human and Applied Physiological Sciences, School of Basic and Medical Biosciences, Faculty of Life Sciences & Medicine, King’s College London, UK; Department of Biomedical Sciences, Faculty of Health and Medical Sciences, University of Copenhagen, Copenhagen, Denmark; Novo Nordisk Foundation Centre for Basic Metabolic Research, Faculty of Health and Medical Sciences, University of Copenhagen, Copenhagen, Denmark; The Folkhälsan Research Center, Biomedicum Helsinki, Helsinki, Finland; Department of Medical Genetics, Medicum, Biomedicum Helsinki, University of Helsinki, Helsinki, Finland; Department of Neurology, Faculdade de Medicina (FMUSP), Universidade de São Paulo, São Paulo, Brazil; Department of Pathology, Medical College of Wisconsin, Milwaukee, WI, USA; Diverge Translational Science Laboratory, Milwaukee, WI, USA; Department of Paediatric Neurology - Neuromuscular Service, Evelina London Children’s Hospital, Guy’s and St Thomas’ Hospitals NHS Foundation Trust, London, UK; Randall Centre for Cell and Molecular Biophysics, Faculty of Life Sciences and Medicine (FoLSM), King’s College London, London, UK

## Abstract

Skeletal muscle is a complex syncytial arrangement of an array of cell types and, in the case of muscle specific cells (myofibers), sub-types. There exists extensive heterogeneity in skeletal muscle functional behaviour and molecular landscape, at the cell composition, myofiber sub-type and intra-myofiber sub-type level. This heterogeneity highlights limitations in currently applied methodological approaches, which has stagnated our understanding of fundamental skeletal muscle biology in both healthy and myopathic contexts. Here, we developed a novel approach that combines a fluorescence based assay for the biophysical examination of the sarcomeric protein, myosin, coupled with same-myofiber high sensitivity proteome profiling, termed Single Myofiber Protein Function-Omics (SMPFO). Successfully applying this approach to healthy human skeletal muscle tissue, we identify the integrate relationship between myofiber functionality and the underlying proteomic landscape that guides divergent, but physiologically important, behaviour in myofiber sub-types. By applying SMPFO to two forms of human nemaline myopathy (*ACTA1* and *TNNT1* mutations), we reveal significant reduction in the divergence of myofiber sub-types, across both biophysical and proteomic behaviour. Collectively, we develop SMPFO as a novel approach to study skeletal muscle with greater specificity, accuracy and resolution then currently applied methods, facilitating that advancement in understanding of SkM tissue in both healthy and diseased states.

## Introduction

Skeletal muscle (SkM) consists of innervated and metabolically active cells (myofibers), organised into syncytial structures that contain the core contractile apparatus, the sarcomere (Frontera & Ochala, 2015). The integrity, function and molecular regulation of SkM are critical for locomotion, respiration and whole body metabolism, and are increasingly recognised as key determinants of both quality of life and life span (Heymsfield, 2024). In genetic SkM disease (myopathies) however, molecular dysregulation impairs SkM function and condition, significantly impacting health outcomes in humans. Myopathy patients, even with similar genetic backgrounds, exhibit considerable heterogeneity in disease onset, symptom manifestation, progression and life-expectancy. The basis for such variability remains inadequately understood, but the current lack of knowledge is likely to reflect the inherent complexity of SkM tissue.

Human SkM is a heterogenous composite of cell types, including myofibers themselves, that exist to maintain homeostasis of the tissue. These primary resident SkM cells, myofibers, exist in a spectrum of sub-types ranging from oxidative, fatigue resistant and ‘slow’ contracting, through to highly glycolytic and ‘fast’ contracting fibers (Schiaffino & Reggiani, 2011). The precise cellular composition and myofiber sub type proportions found within SkM, and, more importantly in the traditionally analysed whole muscle biopsy homogenates, vary significantly depending on internal (e.g. anatomical site) and external (e.g. diseased state) factors, as revealed by recent single cell analyses (e.g. (Kim *et al*, 2020)). Importantly, there exists vast differences in the molecular programmes of myofiber and non-myofiber cells, and, as we have recently highlighted, even within myofiber sub type populations (Moreno-Justicia *et al*, 2023). These observations highlight that even subtle changes in the underlying composition of (non)myofiber cells within whole muscle homogenates used for analyses, will greatly alter the molecular ‘snapshot’ of the tissue, and may lead to false positives in data sets (Seaborne & Ochala, 2023).

Recent advances in single cell approaches have made it possible to overcome cell composition issues within SkM research. Nonetheless, traditional single cell sequencing has two major shortfalls (Seaborne & Ochala, 2023). Firstly, as typical single nuclei approaches use and profile whole sample lysates, they lose the capacity to trace molecular data back to the individual myofiber cell they are derived thus losing myofiber subtype specificity within the data set. While single myofiber sequencing protocols overcome this issue (Blackburn *et al*, 2019; Sahinyan *et al*, 2022), they only yield a single data set per single myofiber, which, given the vast inter-myofiber subtype molecular heterogeneity, falls short at completely profiling single muscle fibers both on molecular and functional level. While methods elsewhere have obtained dual-omic data from the same single cell (providing insights into multi-layers of molecular regulation; (Angermueller *et al*, 2016; Argelaguet *et al*, 2019; Gu *et al*, 2021)), to date, no viable method has been able to develop single cell paired biophysical and omic data in humans, albeit recent work has been performed in fresh rodent tissue (Ng *et al*, 2024). In SkM research, such an approach would allow improved understanding of how underlying myofiber subtype molecular landscapes may impact functional performance of the cell, to a unique level of accuracy and specificity. Combining a functional read out and cell wide global omics from the same single myofiber will not only help to advance our understanding of SkM biology, but also to understand more about the dysregulation that occurs in SkM disease and in the development of myopathic therapeutics more holistically.

Here, we present **<u>S</u>**ingle **<u>M</u>**yofiber **<u>P</u>**rotein **<u>F</u>**unction-**<u>O</u>**mics (SMPFO), a unique single myofiber workflow, enabling examination of the biophysics of the sarcomeric protein, myosin, coupled with global proteomic profiling. We applied this method to interrogate the myofiber sub-type differences in control SkM tissue, and in SkM of two forms of human nemaline myopathy, a genetically diverse congenital myopathy with characteristic abnormalities and defects predominantly in proteins affecting normal sarcomeric assembly and contractile function (Jungbluth *et al*, 2018). Our approach reveals coordinated and physiologically important myofiber sub-type specificity across biophysical and proteome levels, that, in in a SkM diseased state, is largely lost or ‘shrunk’. As SMPFO is universally applicable to study myofibers in all contexts it may provide the basis for a new understanding of SkM biology in health and disease.

## Results

### Single myofiber protein function-omics uniquely correlates myosin biochemical state and global proteome data

We developed a unique single myofiber workflow, enabling examination of the biophysics of the sarcomeric protein, myosin, coupled with sensitive global proteomic profiling - termed **<u>S</u>**ingle **<u>M</u>**yofiber **<u>P</u>**rotein **<u>F</u>**unction-**<u>O</u>**mics (SMPFO) (Fig. 1A). Using this workflow on control SkM biopsies (N=8; Supp. table 1), we successfully resolved both single-myofiber global proteome data (Fig. 1C) and the biochemical state of myosin, i.e., super-relaxed state (SRX; Fig. 1D), in 68/91 original myofibers analysed (∼75%; Fig. 1B). Here, in keeping with previous findings from our group (Carrington *et al*, 2023), we show a highly heterogenous spread of myosin shape in healthy SkM, suggesting that myosin exists in an array of poised states for relaxation or contraction (Fig. 1D).

A major benefit of our single myofiber workflow is in its ability to generate single myofiber proteome data, enabling an antibody-independent myofiber sub-type classification, beyond classical type I (slow) and type II (fast) isoforms. Using highly sensitive proteome analyses, we successfully obtained an average of 781 (± 401) protein hits from N=86 individual myofibers (Fig. 1C; myofibers measuring ∼0.5mm in length), with the most abundant proteins largely specific to key muscle Gene Ontology (GO) terms such as ‘muscle contraction (GO:0006939)’, ‘muscle cell development (GO:0055001)’ and ‘muscle system process (GO:0003012)’ (Fig. 1E-F). Using these data, we applied a myosin heavy chain isoform (MYH7, MYH2 and MYH1) specific expression approach to label each myofiber into specific subtypes, as per our recent work (Lewis *et al*, 2024; Moreno-Justicia *et al*., 2023), which, in keeping with our previous findings, we resolve type I, hybrid I/IIa and type IIa myofibers, with no evidence of pure type IIx myofibers (Fig. 1G). We next grouped type I and type IIa myofibers per sample and performed pseudo-bulk analysis of type I vs type IIa myofibers (Fig. 1H). Unsurprisingly, this comparison revealed the most differentially abundant proteins (differentially abundant proteins; Xiao < 0.05) to belong to both metabolic and sarcomeric GO terms, with ‘transition between fast and slow fiber’, ‘oxidative phosphorylation’ and ‘muscle contraction’ to be some of the most significantly enriched terms (Fig. 1I).

Collectively, we prove that our SMPFO methodology is able to adequately resolve the biochemical/functional behaviour of the key sarcomeric protein, myosin, as well as developing accurate and specific global proteome data from the exact same single myofiber. To our knowledge, this is the first developed methodology able to acquire both proteomic and protein functional data from the exact same cell.

### Myosin biochemical state is associated with metabolic and sarcomeric protein expression in a myofiber sub-type specific manner

The true power of our workflow is in the unique opportunity to pair two separate data types, originating from the exact same myofiber. This enables us, for the very first time, to investigate specific underlying proteome signatures and the association with functional outputs of the myofiber. Using our dual-data set (N=68; Fig. 1B) we first clustered myofibers into either low (≤ 35%) or high (≥65%) SRX myosin clusters (Fig. 1D), before performing pseudo-bulk analyses on these subjects. Interestingly, this analysis suggests that myofibers with a low SRX proportion are significantly (Xiao < 0.05; Fig. 1J) enriched in the expression of slow isoform sarcomeric proteins (e.g. TNNI1, TNNC1, and TNNT1; Fig. 1J). Conversely, myofibers with a high SRX percentage have a greater abundance of proteins associated with important roles in metabolic or ATP activity processes (PFKM, CAMK2B, ACAA2) or with roles in ribosomal biology (RPL24, RPL35a; table). We corroborated these findings by correlating single myofiber SRX with proteomic data from the same myofiber (Figure 1K; unadjusted, see methodology). Here, by overlapping significant hits identified in both our correlation (Figure 1K) and differential abundance analysis (Figure 1J), we found 18 proteins commonly identified. This list of 18 hits highlights a negative association between SRX proportion, and a number of proteins linked with ATPase activity, including VWA8 (R=-0.46, *P*=0.029) and ATP2A2 (R=-0.29, *P*=0.045), suggesting that greater SRX is associated with a reduction in the abundance of proteins with known ATPase roles.

Taking advantage of our myofiber sub-type data set, we deepened our correlation analysis to a myofiber sub-type specific level (figure 1L). Analysing only myofibers with both proteome and SRX data, and only proteins that were identified in 50% of myofibers (per sub-type analysis, see methods), we identify 39 proteins who significantly correlate in at least one of the myofiber sub-types (figure 1L). We show clear myofiber sub-type specificity within these 39 proteins. For example, the protein, Sarcoglycan Delta (SGCD; Figure 1M), displays significant, but divergent associations between SRX and SGCD abundance, in type I (R=0.63, *P*=0.038) and type IIa myofibers (R=-0.55, *P*=0.049) myofibers.

Collectively, SPMFO highlights the heterogeneity, subtype specificity and relationship of myosin biochemical state and underlying protein expression in human SkM. We have previously shown that SkM myopathies display dysregulation in myosin functionality and the underlying proteomic landscape (Laitila *et al*, 2024), but the extent of coordination and myofiber sub-type specificity in this response is unknown.

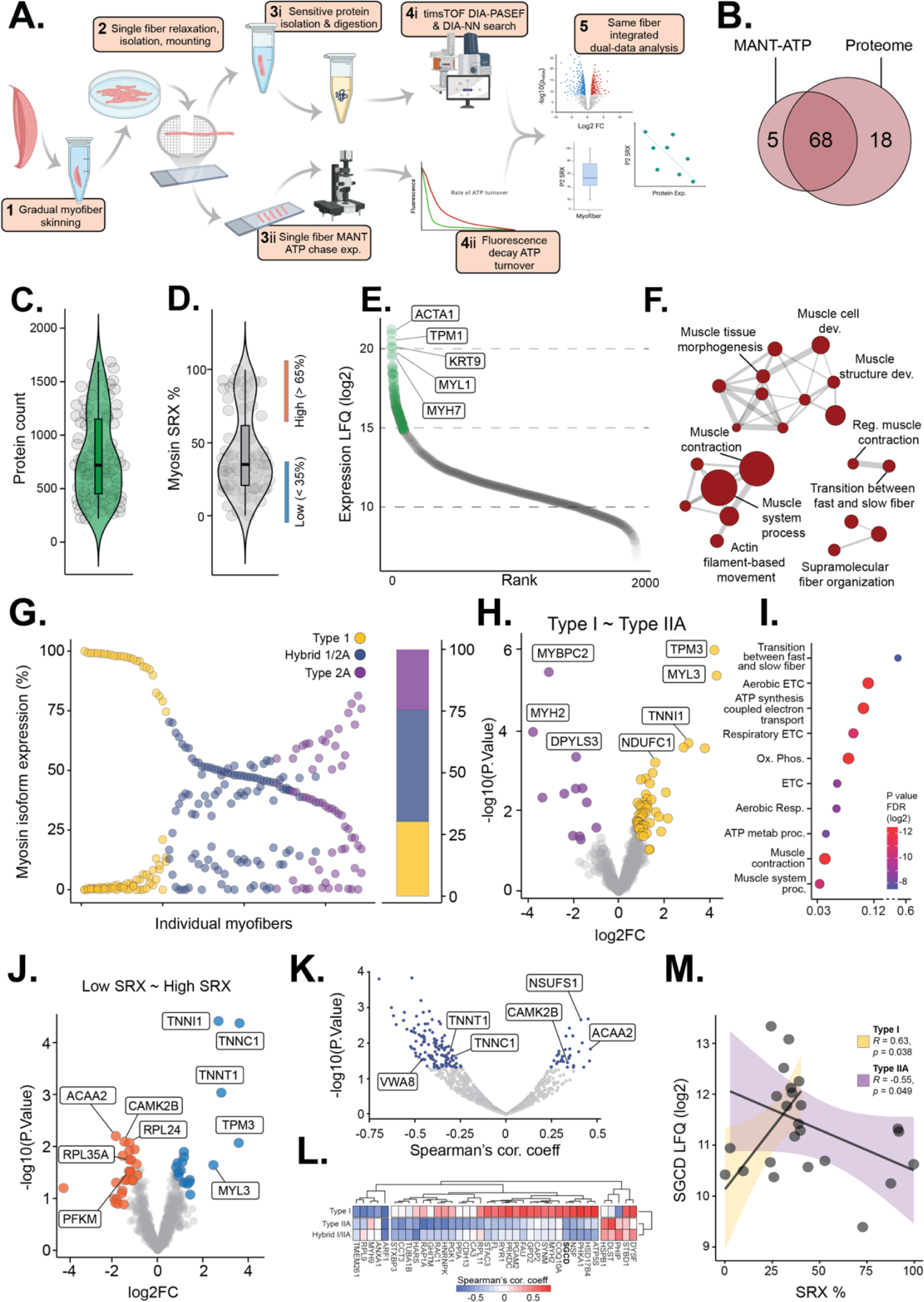

### Nemaline myopathy patients display similarly dysregulated myosin behaviour but diverse myofiber composition

We employed our workflow to patient SkM biopsies derived from Nemaline Myopathy patients with mutations in *ACTA1* (N=3) or *TNNT1* (N=3), and controls (N=3; Supp. table 1) from our original analysis. Despite the manually isolated myofibers of *ACTA1-* and *TNNT1-* samples being visibly smaller than control (Fig. 2A), our SMPFO method was able to successfully develop a dual-data set of N=56 (80% success rate; Fig. 2B). In significant contrast to the array of SRX percentage of myofibers in control SkM myofibers (Con; 54.5 ± 32.0 %; Fig. 2C), myofibers of *ACTA1-* and *TNNT1-* NM patients are more homogenous, with a shift to a lower SRX proportion (*ACTA1-*, 38.4 ± 8.6%, P=0.08; *TNNT1-*, 33.0 ± 14.8 %, P=0.03). This reduction in SRX myosins would suggest an increase in ATP demand and turnover within myofibers of *ACTA1-* and *TNNT1-* SkM (Ranu *et al*, 2022).

Unsurprisingly, the number of identified proteins from our global proteomic analysis was incrementally reduced by disease severity (Con, 1141 ± 331; *ACTA1-*, 729 ± 218; *TNNT1-*, 635 ± 256; Fig. 1D), but GO analysis of the most abundantly identified proteins suggested a continued high degree of SkM specificity (Fig. 1E-F). Using our myosin heavy chain isoform subtype classification approach, we observed a clear shift in myofiber subtype composition, with *ACTA1-* and *TNNT1-* NM muscle containing a greater percentage of type I and hybrid/type IIa myofibers, respectively (Fig. 2G). This observation, supported by immunohistochemistry analyses of control and NM myofibers (Fig. 2A), is an interesting observation given that these patients classically present with similar clinical pathology..

### The oxidative metabolic proteome is commonly dysregulated across Nemaline Myopathy conditions, despite the differing myofiber subtype compositions

To develop a deeper understanding of what may underpin myofiber subtype divergence, but similar behaviour in myosin biochemical state, we performed differential proteomic analysis across specific myofiber subtypes in NM and control samples (Fig. 2H & 2I). We observed a large disruption in the proteomes of both *ACTA1-* type I (158 differentially abundant proteins) and *TNNT1-* type IIa myofibers (176 differentially abundant proteins), compared to the same control subtypes. When we analysed only the statistically down-regulated proteins in these comparisons, we found surprising commonality in the enrichment of GO terms (Fig. 1J). Notably, ‘aerobic respiration’ (GO:0009060) was the most significantly enriched term across both *ACTA1-* type I and *TNNT1-* type IIa analyses, as well as being one of the top 10 most enriched terms in our original control type I ∼ type IIa myofiber comparison (Fig. 1I). This suggests that, irrespective of present myofiber sub-types, NM induces a ubiquitous dysregulation of the energy and oxidative metabolism-related proteome in NM SkM. We therefore investigated the extent of this synergistic dysregulation across type I (*ACTA1-*) and type IIa (*TNNT1-*) myofibers in NM, versus control. We overlapped the differentially abundant proteins from these original differential expression comparisons (e.g. Fig. 2H and Fig. 2I), identifying 79 common differentially abundant proteins across NM disease types (Fig. 2K). Here, an overwhelming proportion of these proteins (78/79) displayed common directionality in dysregulation, including the down-regulation of key metabolic proteins such as COX5A, PYGM and PDHA1 (Fig. 2L).

### The diversity of the myofiber subtype proteome and the relationship with myosin protein function, is lost or ‘shrunk’ in NM

The only exception to the trend in common directional changes in these 79 proteins was MYBPC2 (Fig. 2L). Indeed, MYBPC2 was the only protein significantly dysregulated in both comparisons, but with different directionality of change (Fig. 2L). MYBPC2 as the ‘fast’ paralog of the MYBPC protein family is traditionally more enriched in type II myofibers. Indeed, we found MYBPC2 to be one of the most differentially abundant proteins in our control data set (type I ∼ type IIa; Fig. 1H) and confirmed this in our disease subset data (Fig. 2M). However, this myofiber subtype specific abundance pattern, is dysregulated in NM SkM. Indeed, MYBPC2 abundance is increased in *ACTA1-* type I myofibers (14.15 ± 2.27) compared to control type I (P = 0.067), and significantly decreased in *TNNT1-* type IIa (15.04 ± 1.18) vs control type IIa myofibers (P < 0.05), resulting in a non-significant difference between the two NM myofibers sub-types (*ACTA1-* type I ∼ *TNNT1-* type IIa, P = NS; Fig. 2M). These data suggest that the myofiber subtype specificity in expression of the fast paralog of the MYBPC family, is lost or ‘shrunk’ in NM.

We sought to ascertain how extensive this ‘shrinking’ in myofiber subtype diversity in NM SkM is. We theorised that by comparing differentially abundant proteins within our NM myopathy subtype myofibers (e.g. *ACTA1-* type I ∼ *TNNT1-* type IIa) against those found in a control type I vs type IIa data set, we would be able to inspect the degree of maintenance of myofiber subtype diversity across control to NM disease. We performed differential abundance analysis in our matched control (control type I ∼ control type IIa, 78 significant hits, N=3) and NM disease samples (*ACTA1-* type I ∼ *TNNT1-* type IIa, 50 significant hits, N=3), where 35 proteins from our control differentially abundant proteins list that were identified in the entire NM data set (Fig. 2N). Of these 35 control differentially abundant proteins, the abundance of 21 proteins was non-significant between *ACTA1-* type I and *TNNT1-* type IIa myofibers (Fig. 2N), suggesting a loss myofiber subtype specificity. A large proportion of these proteins have important roles in mitochondrial function (CKMT2, COA3, COQ9, UQCRC1), energy metabolism/processing (ATPA2, ATP5J, PYGM) and myofiber/sarcomere structure and function (ACTN3, MYBPC2, MYH1, MYOM3, MYOZ1, TUBB). The remaining 14 proteins, largely sarcomere isoform specific proteins including MYH2, MYL1/2/3, TNNC1/2, TNNT1/2 and TPM3, maintained significance between myofiber subtypes in NM disease (*ACTA1-* type I vs *TNNT1-* type IIa). However, by correlating the differences between myofiber subtypes within each comparison (e.g. control type I ∼ type IIa vs *ACTA1-* type I ∼ *TNNT1-* type IIa; Fig. 2O), we observed a reduction in fold-change in NM disease. For example, the difference in abundance of MYL2 is markedly greater in our control type I vs type IIa analysis, then in the NM comparisons (Fig. 2O). This trend is observed across 12 of the 14 significant differentially abundant proteins.

Taken together, our work, using a novel single myofiber methodology, identifies important homeostatic divergence between type I and type IIa myofibers in control SkM tissue, that in SkM of patients with NM, is greatly reduced.

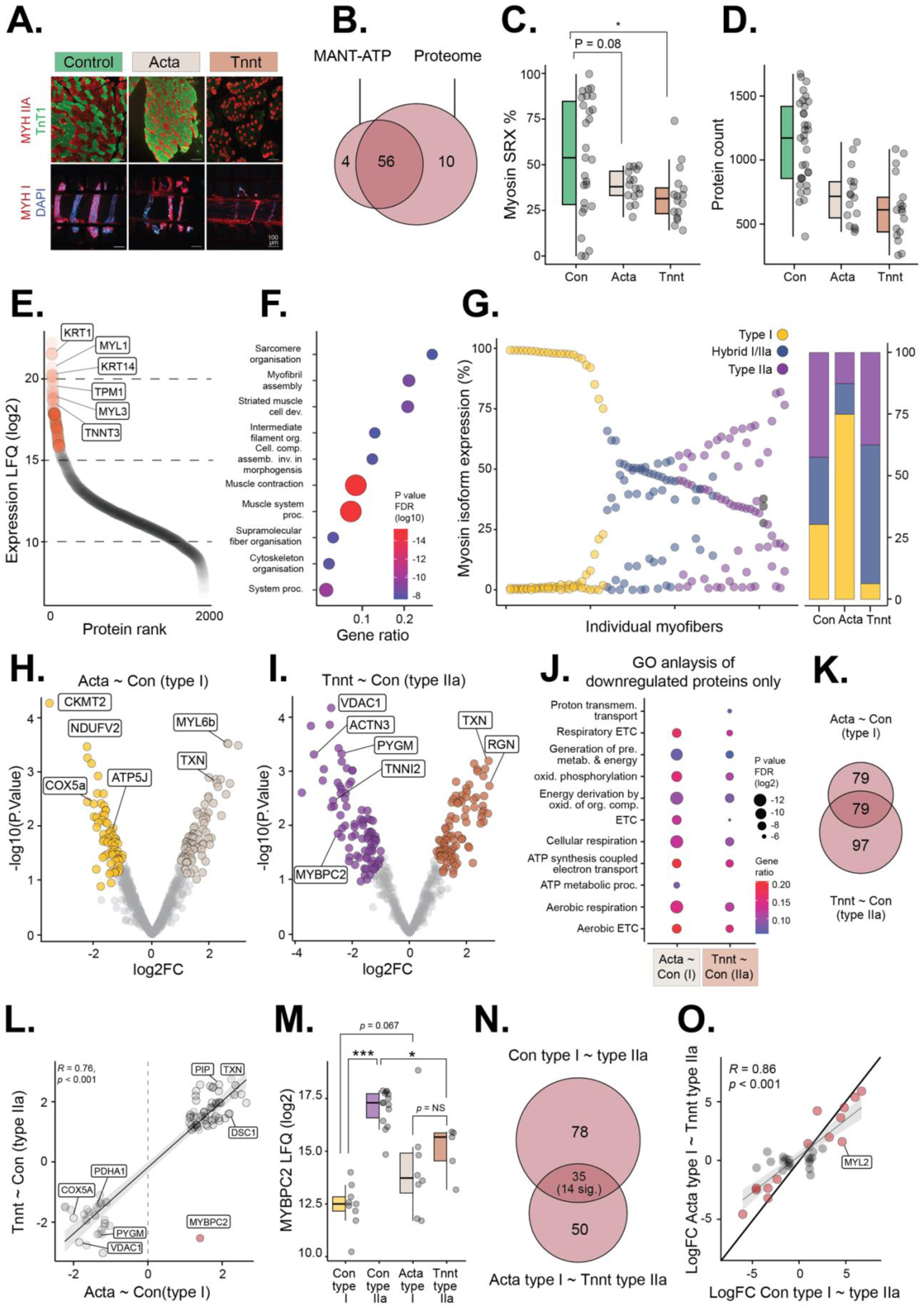

## Discussion and summary

Human skeletal muscle is a highly heterogenous composition of different cell types and sub cell types, archetypally observed in the diverse spectrum of myofiber types (Schiaffino & Reggiani, 2011). This complex cellular arrangement is crucial for supporting homeostatic tissue function, but, as we have recently discussed (Seaborne & Ochala, 2023), presents complications when analysing SkM in healthy and diseased contexts. Here, to the best of our knowledge, we present the first methodology that enables profiling of single protein biophysical function and global protein expression, originating from the exact same single cell (myofiber) from human SkM. This approach identifies important myofiber subtype coordination between protein abundance and myosin biophysical behaviour in control SkM tissue. Our method shows that such diversity in protein function and abundance is lost, or shrunk, in myofibers of NM patients. Collectively highlighting the unique power of SMPFO to transform our understanding of human SkM in both healthy and diseased contexts and to support the advance of therapeutics and interventions to alleviate SkM disorders.

The most important and unique aspect of SMPFO is in its ability to resolve functional and protein expression data originating from the same myofiber (termed protein function-omics), allowing highly accurate associations between data sets. Categorising single myofibers into either high or low myosin biophysical shape, based upon SRX percentage), we identify a signature of protein expression that associates with myosin SRX. The percentage of myofiber SRX is an important variable in SkM tissue and whole body metabolic demands. Indeed, it is purported that if all myofibers switched from the energy conservative SRX state, to the more ATP demanding disordered relaxed state (DRX), this would increase whole body metabolism by ∼50% (Wilson *et al*, 2021). We show here, for the very first time, that myofibers in a high SRX state have an increased abundance of metabolic related proteins, including PFKM, CAMK2B and ACAA2, whereas myofibers with a reduced SRX (and concomitant increased DRX) state, are enriched for key sarcomeric proteins (TNNI1, TPM3, MYL3). Single myofiber protein-function omic correlative analysis further identifies a negative association between proteins involved in ATP activity (VWA8 and ATP2A2) and the percentage of myosin SRX. Collectively, these data suggest that, in control SkM tissue, myofibers that are in a quiescent, energy-replenished state have an increase in metabolically related proteins and those in a disordered, energy-consuming state show an increase in proteins that are involved in ATP turnover and sarcomeric organisation.

Single cell omics technologies have revolutionised our approach to studying SkM, revealing the cell-type diversity, complexity and transcriptomic profiles that control and regulate the tissue (Dos Santos *et al*, 2020; Kim *et al*., 2020; Petrany *et al*, 2020). However, due to the underlying chemistry, typical single cell approaches lose the capacity to trace data back to individual myofiber subtypes and cannot yield combined functional and molecular data sets from the same myofiber. By analysing myofibers on an individual basis, SMPFO overcomes these shortfalls thereby supporting analysis of protein function-omic data on a myofiber specific subtype basis. Here, we correlate myosin SRX percentage with protein expression in type I and type IIa control myofibers, identifying divergent associations in array of proteins. Interestingly, we reveal the correlation between SGCD and SRX percentage, that is significantly divergent depending on the specific myofiber subtype analysed. SGCD, a component of the Sarcoglycan complex and dystrophic-glycoprotein complex, is involved in linking F-actin cytoskeleton to the extracellular matrix with mutations in the coding gene leading to muscular dystrophy and dilated cardiomyopathy (Coral-Vazquez *et al*, 1999; Nigro *et al*, 1996). The myofiber subtype specific association with SRX percentage is an interesting and unique observation, as very little is known or reported regarding SGCDs potential role in energy consumption in SkM. Thus further work is needed to validate and expanding this finding. Collectively, we show that SMPFO is able to decipher clear myofiber subtype specificity and diversity across protein function-omic data in control SkM. Such diversity in myofiber behaviour (functional and molecular) is crucial to meet the demands of life, supporting homeostatic regulation of the tissue and promoting health and prolong life-span in humans (Bottinelli & Reggiani, 2000).

We therefore employed our method to a diseased SkM context, where there is a dearth in knowledge of molecular pathophysiology, to see if we were able to decrypt some of this obscurity. Herein, nemaline myopathy, a sub-class of congenital SkM disease, is an archetypal case (Laitila & Wallgren-Pettersson, 2021). In humans, mutations in at least 12 causal genes predominantly implicated in sarcomeric assembly and function, have been identified leading to a wide spectrum of disease pathology and severity. Histologically, it has routinely been observed that patients with pathogenic mutations in Actin and Troponin present with divergent compositions of myofiber subtypes (enriched for type I and type IIa, respectively), despite presenting with similar disease outcomes (Malfatti & Romero, 2016). Our data is in accordance with these histological observations, and also in support of our recent work on the disease (Moreno-Justicia *et al*., 2023). Interestingly however, despite the diversity in myofiber subtype prevalence, we observed relatively similar disruptions in the proteome of these samples. Indeed, we report a significant, and ubiquitous, reduction in the abundance of proteins relating to metabolic, oxidative and energy relating terms, that in control SkM, underpins a proportion of the diversity between type I and type IIa myofiber subtypes. We also show a clear and significant shrinking of myofiber biophysical behaviour in both forms of NM, irrespective of the underlying myofiber subtype composition. We, and others, have regularly reported that genetic striated muscle disease is associated with a dysregulation in myosin biochemical state (for example, (Anderson *et al*, 2018; Carrington *et al*., 2023)) and significant disruption of proteins relating to metabolism and energy handling (Laitila *et al*., 2024; Ranu *et al*., 2022; Slick *et al*, 2023; Tinklenberg *et al*, 2023), but this is the first report illustrating this data originating from the exact same myofiber.

When examining the uniformity between NM type I and type IIa proteomes, we identified MYBPC2 expression to be counterintuitively modified. That is, we report an increase in *ACTA1-* Type I, and reduction in *TNNT1-* Type IIa MYBPC2, compared to relevant control myofibers, respectively. MYBPC2 is a key regulator of muscle functionality. Indeed, it has been recently reported that homozygous knockout of MYBPC2 induces SkM tissue reductions in grip and plantar flexor muscle strength, as well as speed of contraction and peak isometric force in isolated EDL muscle of mice (Song *et al*, 2021). Our finding of a reduction in MYBPC2 expression across specific myofiber subtype in NM, is mirrored by a reduction in heterogeneity of myosin SRX state, within these same myofibers. There is growing interest in the association between MYBPC and myosin conformation (Lewis & Ochala, 2023). Indeed, mice with either homozygous or heterozygous truncating mutations of the cardiac specific MYBPC paralog (c-MYBPC) report significantly elevated proportion of myosins in DRX state (Toepfer *et al*, 2019). Further work shows that not just the presence, but the post-translational state of c-MYBPC (hyperphosphorylated), also induces an increase in myosin DRX state (Jiang *et al*, 2015), as the knock out models. However, we are the first to report that a reduction in MYBPC2 differential between type I and type IIa myofibers in NM SkM is paralleled by a reduction in myosin conformational shape heterogeneity. This loss of homeostatic myofiber specific divergence across protein function-omic data, and the impact this has on human SkM disease, requires further examination.

This work is not without limitation. Chiefly, the sample size used within this proof-of-principle work, are of low power both at whole biopsy sample and individual myofibers level. The primary findings from this work require further validation through independent analysis of healthy control and nemaline myopathy patient cohort samples. Furthermore, the protocol described herein first uses a graded glycerol approach to skin myofibers, before performing the dual function-omic assay. Chemical skinning of myofibers is necessary for inspection of myosin biochemical conformation (Stewart *et al*, 2010) but likely impacts the quantity of proteomic data we were able to retrieve. While it has recently been suggested that chemical skinning alters structural properties of myofibers (Lewalle *et al*, 2022), to the best of the authors’ knowledge, no work has examined the differential proteome content of fresh ∼ skinned myofibers. Nonetheless, we report a marked reduction in the number of quantified proteins in these samples, compared to our recent work on the same sample set in non-skinned conditions (Moreno-Justicia *et al*., 2023). This may reflect a reduction in sarcolemma and further membrane bound proteins (Lewalle *et al*., 2022). To overcome this, post-processing chemical skinning by Triton-X100 will enable proteomic analyses to be performed on a non-skinned fraction of the single myofiber. The reduction in quantified targets is also likely due to the portioning of a single myofiber into two constituent parts, each for separate analyses, thus reducing the material for single myofiber proteomic analyses. This latter consideration also implicates sub-myofiber compartmentalisation of molecular landscapes. That is, myofibers may possess certain differential molecular domains (e.g. myotendinous junction (Karlsen *et al*, 2023)) along the length of the cell, where our myofiber portioning workflow may impact the myofiber type specificity of our dual-data. Contrary to this hypothesis however, myofibers seems to possess synchronised and coordinated expression of myosin isoforms (Dos Santos *et al*., 2020), thus sub-type is maintained across the entire myofiber cell. Nonetheless, future work would benefit from obtaining all data from the exact same myofiber segment to avoid the confounding of potential molecular compartments within single myofibers.

In summary, we have developed the first methodology that allows simultaneous examination of the biophysical functionality of myosin and the global proteome of the exact same single myofiber. This approach provides unique insights into the important myofiber subtype diversity from both a functional and molecular perspective, within control skeletal muscle, and the extent to which this is dysregulated, and myofiber subtype diversity shrunk, in human nemaline myopathy. Single myofibre protein function-omics could therefore become a pivotal tool in not only extending our understanding of fundamental myology and SkM myopathy, but also for identifying biomarkers and for providing the bases for future therapeutic developments to alleviate SkM disease.

### Methodology

#### Samples collection and processing

The samples for this work were obtained from previous studies (outlined where applicable). For our control cohort, N=8 patient biopsies were obtained from the *vastus lateralis* muscle under local anaesthesia. Samples were immediately snap frozen in liquid nitrogen and stored for further processing (Sahl *et al*, 2018; Søndergård *et al*, 2021). These samples were approved by the ethical committee for the Capital Region of Denmark (H-15010122) and by the local ethics committee of Copenhagen & Frederiksberg (H-15002266), with full participant consent. Six patients with confirmed cases of nemaline myopathy (N=3 *ACTA1-*, N=3 *TNNT1-*), were selected for analysis in our disease study, alongside N=3 control samples from our original control cohort (see Supp. Table 1 for characteristics). The nemaline myopathy patient biopsies were collected, stored and processed in accordance with the Human Tissue Act under local ethical in United Kingdom (REC 13/NE/0373), with consent. All procedures were carried out in accordance with the Declaration of Helsinki. From all samples, a section of the original biopsy was dissected and isolated under sterile, frozen conditions and prepared for processing.

#### Solutions and buffers

For Mant-ATP chase experiments: relaxing solution contained 4 mM Mg-ATP, 1 mM free Mg2+, 10-6 mM free Ca2+, 20 mM imidazole, 7 mM EGTA, 14.5 mM creatine phosphate and KCl to an ionic strength of 180 mM and pH of 7.0. Rigor buffer contained 120 mM K acetate, 5 mM Mg acetate, 2.5 mM K2HPO4, 50 mM MPS and 2 mM DTT with a pH of 6.8.

Proteomics: lysis buffer contained 1% sodium dodecyl sulfate, 40 mM chloroacetamide, 10 mM dithiothreitol in 50 mM Tris with a pH of 8.5.

#### Equations

Double exponential Mant-ATP decay:

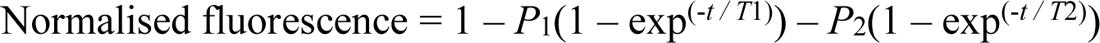

Note: *P_1_ = amplitude of the initial decay; T_1_ = time constant for P_1_; P_2_ = amplitude of the secondary decay; T_2_ = time constant for P_2_*

### Single fiber dual-assay processing

Individual myofibers were then transferred to microscopy slides containing single half-split copper meshes designed for electron microscopy (G100 2010C:XA; SPI Supplies, West Chester, PA, USA; width, 3 mm). Myofibers were clamped under copper grids with N of 5/6 per copper mesh, per microscopy slide. Once clamped, the end tail of each individual myofiber was dissected and this micro-piece of myofiber was transferred, via micro-tweezers, to a single well of a 96-well plate on ice, containing proteomic lysis buffer (15ul; see above). Between the handling and dissection of each individual myofiber, implements (micro-tweezers, scalpel, syringe needles) were washed through with RNAse zap, 100% EtOH and left to air-dry. Multiple sets of implements were used during experiments to ensure efficient and fast processing of all myofibers. Herein, each single myofiber pertained two micro-fragments, one piece for MANT-ATP chase experiments, and one piece for single myofiber global proteomic profiling (Fig. 1A).

### MANT-ATP chase assay

Single myofibers were mounted in accordance with our previous studies (Laitila *et al*., 2024; Lewis *et al*., 2024; Ochala *et al*, 2021). Cover slips were placed to the top of the microscopy slide, held with double-sided tape to generate ‘flow’ chambers where myofibers were incubated in rigor buffer for 5-mins at 25°C. A rigor buffer containing 250 um Mant-ATP was then flushed through the chamber and left to incubate the myofibers for 5-mins. A secondary rigor buffer containing unlabelled ATP (4Mm) was flushed into the chamber, and acquisition of the Mant-ATP chase occurred.

### MANT-ATP fluorescence acquisition

To acquire fluorescence decay in individual myofibers, a microscopy setup containing AxioCam ICm1 camera (Zeiss) with Plan-Apochromat 20X/0.8 objective and Axio Scope A1 microscope (Zeiss), took images (20 ms exposure time using a DAPI filter set) every 5 s for the first 90 s and every 10 s for the remaining 5 minute acquisition time. From each myofiber, three independent regions of the same myofiber were samples for fluorescence decay using region of interest (ROI) manager (ImageJ; Bethesda, MD, USA). A normalised mean fluorescence intensity value was generated by taking the background intensity away from each individual image, averaged across the three images and normalised to the final Mant-ATP image (T = 0). Data was exported to Prism (V9.0; GraphPad Software Inc., San Diego, CA, USA) and fitted to an unconstrained double exponential decay (see equations). The double exponential decay indicates the initial rapid decay (P1) as percentage of myosin in DRX state, and the slower, secondary decay (P2) as the percentage of myosin in the SRX state (Ochala *et al*., 2021).

### Myofiber proteomic sample preparation

Single muscle fiber proteomics was performed as recently described (Moreno-Justicia et al., 2023). In brief, proteomics lysis buffer was added to each myofiber piece and boiled at 95℃ in a thermomixer before sonication with a bioruptor instrument. Myofiber lysates underwent protein digestion by addition of 1:100 trypsin (Promega) and 1:500 LysC (Wako), the reaction was allowed to take place overnight in a thermomixer set at 37℃. The next day, protein digestion was stopped by addition of isopropanol 2% trifluoroacetic acid and the tryptic peptides were desalted using in-house crafted styrenedivinylbenzene reverse-phase sulfonate (SDB-RPS) stage-tip columns. Lastly, desalted peptides were loaded in Evotips (Evosep) prior LC-MS/MS analysis following the company’s loading protocols.

### Liquid chromatography tandem mass spectrometry

The LC-MS/MS instrumentation consisted of an Evosep One HPLC system (Evosep) (Bache *et al*, 2018)coupled via electrospray ionization to a timsTOF SCP mass spectrometer (Bruker). Peptides were separated using an 8 cm, 150 μM ID column with C18 beads (1.5 μm). Chromatographic separation followed the “60 samples per day” method, and electrospray ionization was performed via a CaptiveSpray ion source. Single muscle fiber peptides were measured using DIA-PASEF (Meier *et al*, 2020), with a scan range of 400-1000 m/z. The TIMS mobility range was set to 0.64-1.37, and cycle time was 0.95 seconds, using 8 DIA-PASEF scans.

### Data processing

Raw MS spectra were processed using the DIA-NN software (version 1.8) (Demichev *et al*, 2020). The search was conducted in a library-based manner, utilizing a previously developed myofiber-specific library (Moreno-Justicia *et al*., 2023). Proteotypic peptides were selected for quantification and the neural network was operated in double-pass mode. Robust LC (high accuracy) was the quantification strategy of choice, together with the match between runs option and a precursor false discovery rate of 1%. Unless specified, other DIA-NN settings remained as default. Downstream bioinformatics were conducted using the PG_matrix file from the DIA-NN output. Principal component analysis was performed using Log2 transformed LFQ values for all myofibers, using *prcomp()* omitting NA values. This revealed one control myofiber (sample; C7, type; I/IIa hybrid) to be an outlier, that was removed. Fiber typing of individual myofibers for both control and NM cohorts was performed as previously described (Moreno-Justicia *et al*., 2023), with small modifications. Briefly, on a single myofiber basis, the raw LFQ values for three myosin isoforms (MYH1, MYH2 and MYH7) were used to calculate their respective relative expression. Samples were then ordered by MYH7 from high to low, and an absolute threshold in expression was used to assign each myofiber to a specific sub type (type I, type IIa, hybrid I/IIa).

### Differential expression

Pseudobulk differential expression analysis was performed by mathematically down sampling each participants total data to create one median data value, per protein hit, per MYH-based fiber type. The limma (v.3.54.2) workflow was followed, including a quantile normalisation of all samples and differential expression (linear model) analysis across fiber sub-types in control cohort, and across conditions in the NM cohort. The Xiao significance score was applied to identify significant proteins (< 0.05), which considers the fold change and statistical significance of protein hits (Xiao *et al*, 2014). To note, for SRX % differential proteome analysis, N=7 samples contained both low (≤ 35%) and high (≥ 65%) SRX myofiber sub-types and were used for analysis.

### Functional enrichment

Gene ontology functional enrichment analysis was performed focussing on ‘biological process’ terminologies, unless otherwise stated. List of relevant protein hits (identified within relevant analyses) were exported to ShinyGO (v.0.77 or v.080; (Ge *et al*, 2020)), with the entire data set or comparative data set used as associated background, to ascertain functional enrichment terms with an adjusted FDR cutoff of < 0.05 interpreted as significant. Gene ratio was ascertained by dividing number of identified genes in the input list, by the total number of genes within the term. Where relevant, packages REVIGO (v.1.8.1 (Supek *et al*, 2011)), MetaScape (Zhou *et al*, 2019) and Cytoscape (v.3.9.1) were used to develop and represent GO association networks.

### Correlation coefficient and hierarchical clustering

Data was filtered to remove outlier (N=1) and non-dual data (N=23) samples, and protein data present in < 22 of these samples, retaining a data set of N=67 samples and N=979 protein hits. Log2 LFQ protein values were correlated (Spearman, unadjusted) against SRX and DRX % in remaining samples via *rcorr()* function in Hmisc package (v 5.1-3). Myofiber subtype correlation analysis was performed as identical to that above, with the exception that protein hits must be present in ≥ 10 samples of each myofiber subtype. Hierarchical clustering was performed on significant type I and type IIa ∼ SRX correlations, using *pheatmap()* function from the Pheatmap package (v 1.0.12). Clusters were determined by visual inspection performed via *cuttree()* function.

### Immunolabelling of muscle sections

Cryosections (10 um) were fixed in 4% PFA (10 mins), permeabilised with Triton X-100 (0.1% for 20 mins) and blocked in 10% Normal Goat Serum (500627, Life Technologies) supplemented with 0.1% BSA (1 h). Sections were incubated overnight with diluted (1:25) primary antibodies against MYH7 (mouse monoclonal A4.951, Santra Cruz, sc-53090) or MYH2 (mouse monoclonal SC71, DSHB), combined with a further antibody against TNNT1 (1:500; rabbit polyclonal HPA058448, Sigma) in 5% goat serum (supplemented with 0.1% of BSA and Triton X-100). Alexa Fluor Goat anti-Mouse 647 (A21237) and Alexa Fluor Dinkey anti-Rabbit 448 (A11034) were used for primary MYH and TNNT1 antibodies, respectively (1:500 dilution in 10% Normal Goat Serum). A fluorescence microscopy setup (10x objective, Zeiss Axio Observer 4 fluorescent microscope, Colibri 5 led detector, Zeiss Axiocam 705 mono camera, Zen Software, Zeiss) captured images of all conditions. A selection of myofibers were mounted on copper grids and imaged through a stereomicroscope, for visualisation purposes. Fluorescent images were obtained with a 10x objective on a Zeiss Axio Observer 3 fluorescence microscope with a Colibri 5 led detector, combined with Zeiss Axiocam 705 mono camera, using Zen software (Zeiss). For visualization purposes, a selection of fibres were mounted on copper grids glued on a microscopy slide and imaged under a stereomicroscope.

### Statistical Analysis

All data, unless otherwise stated, are presented as means ± SD and where possible presented with individual data points. For all Mant-ATP experiments, data was prepared in Prism (V9.0), exported to R Studio via .csv format where it was statistically analysed (ANOVA, with Tukey HSD post hoc) to test for significant (p < 0.05) differences between groups/conditions. Log2 transformed LFQ values for individual protein targets were extracted and an ANOVA with Tukey’s post hoc used to compare across groups/conditions (RStudio). Statistical analysis of proteomic data is detailed in relevant sub-sections.

## Acknowledgements

We thank Thomas Nyegaard Beck for his assistance in many of the experiments outlined in the present manuscript. Mass spectrometry was performed by the Proteomics Research Infrastructure (PRI) at the University of Copenhagen (UCPH), supported by the Novo Nordisk Foundation (NNF19SA0059305).

## Author contributions

Conceptualisation; RAES, ASD, JO. Data curation; RAES, RM-J, JL, CTAL, LS, ASD, JO. Formal analysis; RAES, RM-J, JL, CTAL, ASD, JO. Funding acquisition; RAES, ASD, JO. Investigation; RAES, RM-J, JL, CTAL, LS, EZ, MWL, ASD, JO. Methodology; RAES, RM-J, ASD, JO. Project administration; RAES, JO. Visualisation; RAES. Writing – original draft; RAES, JO. Writing – review & editing; RAES, RM-J, JL, CTAL, LS, EZ, MWL, HJ, ASD, JO

## Data availability

All proteomic data will be submitted to the PRoteomics IDEntification Database (PRIDE) to be made publicly available upon formal submission to peer reviewed journal.

## Supplementary

**Supp. Table 1.** Characteristics of human muscle biopsy samples.

**Supp. File 1.** Differentially expressed proteins across comparative analysis in control and NM disease data sets

**Supp. File 2.** Significant enriched gene ontology terms from control and NM disease data sets

**Supp. File 3.** Significant spearman coefficient correlation analyses.

### Supplementary tables

**Supp. Table 1.**
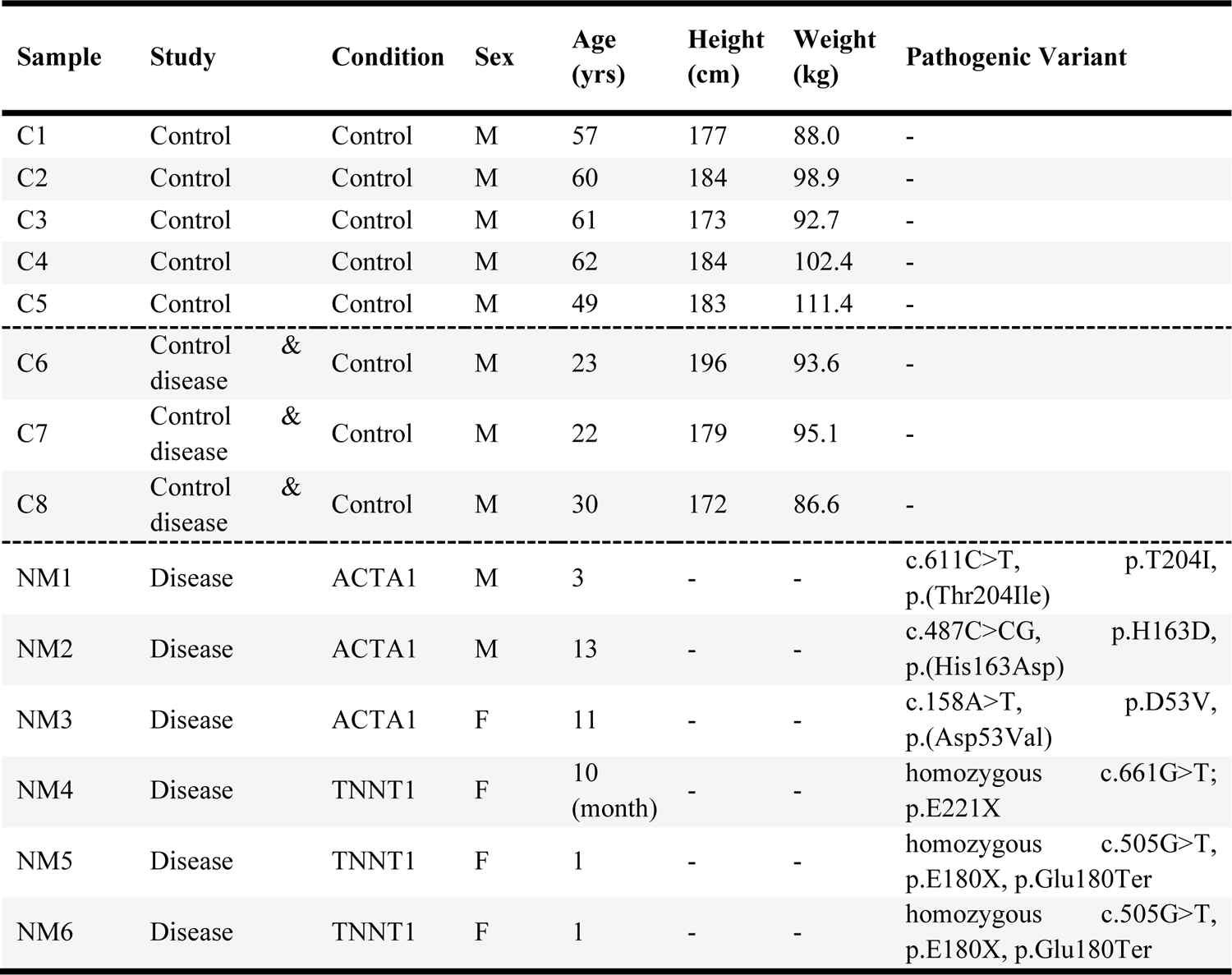
Descriptive of human muscle biopsy samples.

